# *Malassezia sympodialis* Mala s 1 allergen is a potential KELCH protein that cross reacts with human skin

**DOI:** 10.1101/2022.11.02.514932

**Authors:** Dora E. Corzo Leon, Annika Scheynius, Donna M. MacCallum, Carol A. Munro

## Abstract

*Malassezia* yeast species are the dominant commensal fungal species of the human skin microbiota, but are also associated with inflammatory skin diseases, such as seborrheic dermatitis and atopic eczema (AE). Mala s 1, a β-propeller protein, is an allergen identified in *Malassezia sympodialis* inducing both IgE and T-cell reactivity in the majority of patients with AE. In this study, we aimed to elucidate the role of Mala s 1 allergen in skin disease. An anti-Mala s 1 antibody was used to investigate the cellular localisation of Mala s 1, the potential of Mala s 1 as a therapeutic target and examine cross-reactivity of the anti-Mala s 1 antibody with human skin. We demonstrate by high pressure freezing electron microscopy and immune-staining that Mala s 1 is located in the cell wall of *M. sympodialis* yeast cells. Despite the ability of the anti-Mala s 1 antibody to bind to yeast cells, it did not inhibit *M. sympodialis* growth suggesting Mala s 1 may not be an attractive antifungal target. The Mala s 1 predicted protein sequence was analysed *in silico* and was found to contain a motif indicative of a KELCH protein, a group of β-propeller proteins. Humans express a large number of KELCH proteins including some that are localised in the skin. To test the hypothesis that antibodies against Mala s 1 cross react with human skin proteins we examined the binding of anti-Mala s 1 antibody to human skin explant samples. Reactivity with the antibody was visualised in the epidermal layer of skin. To further characterise putative human targets recognised by the anti-Mala s 1 antibody, proteins were extracted from immunoblot gel bands and proteomic analysis performed. Several candidate human proteins were identified. To conclude, we propose that Mala s 1 is a KELCH-like β-propeller protein with similarity to human skin proteins and Mala s 1 recognition may trigger the production of cross-reactive responses that contribute to skin diseases associated with *M. sympodialis*.

## Introduction

*Malassezia* is an abundant genus of 18 yeast species (Vijaya Chandra *et al*, 2021) that includes the dominant commensal fungi of the human skin microbiota (Findley *et al*, 2013), with most prevalent distribution in skin areas of high sebum enrichment (Byrd *et al*, 2018). Besides being commensal skin colonizing yeasts, *Malassezia* is also linked to different skin diseases such as seborrheic dermatitis and atopic eczema (AE) (Saunders *et al*, 2012) and bloodstream infections in immunosuppressed individuals (Vijaya Chandra *et al*, 2021).

AE is a chronic inflammatory skin disease affecting up to 20% of children and 3% of adults (Nutten, 2015; Brunner *et al*, 2017). Genetic and environmental factors leading to impaired skin barrier function have been associated with the pathogenesis of AE (Nutten, 2015). Several human proteins have roles in maintaining skin barrier function, for example filaggrin and filaggrin-2 are important proteins in barrier formation and skin moisturisation (Wu *et al*, 2009), while epiplakin and desmoplakin are essential in epidermal integrity and wound healing (Vasioukhin *et al*, 2001; Ishikawa *et al*, 2010). Skin barrier impairment results in increased contact with different microbes, such as the commensal yeast *Malassezia*, perpetuating skin damage, as well as inducing local and systemic inflammatory reactions (Nutten, 2015). Ten IgE-binding proteins (Mala s 1 and Mala s 5 to 13) have been characterized in *M. sympodialis* (Gioti *et al*, 2013), one of the most frequent species colonizing the skin of both healthy individuals and AE patients in Europe (Jagielski *et al*, 2014; Falk *et al*, 2005). These proteins act as allergens, inducing both IgE and T-cell reactivity in more than 50% of patients with AE (Scheynius *et al*, 2002; Vilhelmsson *et al*, 2007a; Balaji *et al*, 2011).

One hypothesis linking *M. sympodialis* to the pathogenesis of AE is cross-reactivity generated by *Malassezia*-allergens, which are highly homologous to human proteins. Cross-reactivity between *M. sympodialis* allergens and human proteins has been documented for Mala s 11 and Mala s 13 (Balaji *et al*, 2011; Schmid-Grendelmeier *et al*, 2005; Vilhelmsson *et al*, 2007a). Mala s 11 is predicted to be a manganese superoxide dismutase (MnSOD) (Gioti *et al*, 2013; Vilhelmsson *et al*, 2007a). Up to 36% of AE individuals have high levels of specific IgE against human MnSOD and positive skin tests against this allergen (Schmid-Grendelmeier *et al*, 2005). Human and fungal MnSOD induce proliferation of human peripheral blood mononuclear cells (PBMCs) from sensitised AE patients (Schmid-Grendelmeier *et al*, 2005). In addition, CD4^+^ T-cells from AE individuals, specifically sensitised to the *M. sympodialis* Mala s 13 allergen, were reactive to human thioredoxin (Balaji *et al*, 2011). The high similarity with the corresponding mammalian homologs suggests that autoreactive T cells could contribute to tissue inflammation in AE.

Mala s 1 is a 37 kDa protein predicted to be secreted and, to date, has not been shown to have sequence homology to any specific human protein (Zargari *et al*, 1997; Gioti *et al*, 2013). The crystal structure of Mala s 1 revealed it to be a β-propeller protein that binds lipids such as phosphatidylinositols (Vilhelmsson *et al*, 2007b). Mala s 1 is similar to Tri14, a mycotoxin synthesis protein involved in virulence and plant invasion in *Gibberella zeae* (anamorph *Fusarium graminearum*) (Vilhelmsson *et al*, 2007b). Mala s 1 is mainly localized in the *M. sympodialis* cell wall and in the budding area, as visualized by confocal laser scanning microscopy (Zargari *et al*, 1997). Therefore, Mala s 1 may be important for the replication of *Malassezia* and, thus, a potential therapeutic target.

*M. sympodialis* cells can communicate with host cells, such as PBMCs and keratinocytes, through the release of extracellular nanosized vesicles, designated MalaEx (Gehrmann *et al*, 2011; Johansson *et al*, 2018, Vallhov *et al*, 2020). MalaEx are enriched in Mala s 1 compared with the allergens produced by whole *M. sympodialis* cells (Johansson *et al*, 2018). MalaEx induce a different inflammatory cytokine response in PBMCs from patients with AE sensitized to *M. sympodialis* compared to healthy controls (Gehrmann *et al*, 2011), supporting the link between AE and *M. sympodialis*. Recently, the presence of small RNAs in MalaEx was identified, suggesting that they have the potential to deliver functional mRNAs and microRNA-like RNAs to recipient host cells, thereby interfering with the host RNAi machinery to silence host immune genes and cause infection (Rayner *et al*, 2017).

Another potential mechanism of communication with host cells is related to the β-propeller protein structure of Mala s 1 (Vilhelmsson 2017). β-propeller proteins contain different motifs allowing their classification into groups or families (Kopec & Lupas, 2013). One of these β-propeller families contains a KELCH motif (Kopec & Lupas, 2013). The KELCH motif is commonly found in bacterial and eukaryotic proteins with diverse enzymatic functions, and is composed of 50 amino acids, which fold into four stranded beta sheets (Interpro: IPR015915). The interest in KELCH proteins has increased in recent years as these β-propeller proteins are involved in multiple cellular functions and proteinprotein interactions with a wide cellular distribution in intracellular compartments, cell surface and extracellular milieu (Adams *et al*, 2000).

In this study, the aims were to characterise Mala s 1 to a) examine its precise cellular localisation in *M. sympodialis* using high pressure freezing Transmission Electron Microscopy (TEM), b) investigate its potential as a drug target by examining if an anti-Mala s 1 antibody interferes with *M. sympodialis* growth, and c) evaluate potential cross-reactivity between Mala s 1 and human skin proteins, including KELCH proteins, using an anti-Mala s 1 antibody in a human skin explant model.

## Materials and Methods

### M. sympodialis and C. albicans *culture conditions*

*Malassezia sympodialis* (ATCC 42132) was used for all experiments. *M. sympodialis* was grown fresh from −70 °C glycerol stocks for each experiment by plating on Dixon agar containing 3.6% w/v malt extract, 2% w/v desiccated ox-bile, 0.6% w/v bacto tryptone, 0.2% v/v oleic acid, 1% v/v Tween 40, 0.2% v/v glycerol, 2% w/v bacto agar (mDixon) at 35°C for 4-5 days. For culture in mDixon broth (bacto agar omitted), one colony was selected from an agar plate, inoculated into 10 ml of the mDixon culture medium and incubated at 37°C for 3 days in a shaking incubator at 200 rpm. The culture was centrifuged at 200 x g, supernatant discarded, cell pellet washed twice with phosphate buffered saline solution (PBS) and resuspended in 1 ml PBS.

*C. albicans* (ATCC 90028) was grown fresh from −70°C glycerol stocks and cultured in YPD (1% w/v Yeast Extract, 2% w/v Mycological Peptone, 2% w/v glucose) broth overnight at 30°C, shaken at 200 rpm.

### Sample processing and high pressure freezing (HPF) Transmission Electron Microscopy (TEM) imaging

*M. sympodialis* yeasts were cultured as described above on mDixon agar plates, scraped off using a loop and suspended in a minimal amount of distilled water to form a paste. Approximately 2 μl of each sample was processed by HPF using a Leica EM PACT 2 (Leica, Milton Keynes, UK). After HPF, samples were transferred to a Leica AFS 2 embedding system for freeze substitution in a solution with 2% OsO4 in 100% acetone for 40 min. Samples were then placed in 10% Spurr resin:acetone (TAAB Laboratories, Berks, UK) for 72 h followed by 30% Spurr resin overnight, 8 h of 50% Spurr resin, 12 h of 70% Spurr resin, 90% Spurr resin for 8 h and finally embedded in 100% Spurr resin at 60°C for at least 24 h. Sections (90 μm) were cut with a diamond knife onto nickel grids using a Leica UC6 ultramicrotome. Sections were contrast stained with UranyLess EM stain (TABB Group, New York, US) and lead citrate in a Leica AC20 automatic contrasting instrument. Samples were imaged using a JEM 1400 plus transmission electron microscope with AMT ultraVUE camera (JEOL, Welwyn Garden City, UK). These experiments were performed on three biological replicates.

### Gold immunolabelling

Gold immunolabelling was carried out before the contrast staining step described above. Samples on nickel grids were incubated in blocking buffer (1% w/v bovine serum albumin (BSA) and 0.5% w/v Tween 80 in PBS) for 20 min. Samples were then incubated three times for five min in an incubation buffer (0.1% w/v BSA in PBS), then incubated in 5 μg/ml primary antibody in incubation buffer (anti-Mala s 1 mouse monoclonal IgG1 antibody (9G9) from Karolinska Institute (Zagari A *et al*, 1994, Schmidt *et al*, 1997) or mouse monoclonal IgG1 antibody (Agilent Dako X093101-2; Santa Clara, USA) as an isotype control for 90 min. Samples were washed in incubation buffer six times for five min. The secondary donkey-anti-mouse IgG (H&L), Aurion 10 nm gold antibody (810.322; Aurion CliniSciences, Nanterre, France) was added to samples for 60 min (1:40 dilution). Samples were washed six times for five min in incubation buffer. Samples were then washed three times in PBS for five min. Finally, samples were washed in double deionised, filtered water: three times for five min. After these steps, contrast staining was performed as described above.

### M. sympodialis *growth curves and determination of minimal inhibitory concentrations (MICs) in broth microdilution testing*

Growth curve and determination of MICs for voriconazole (Pfizer, Surrey, UK), amphotericin B deoxycholate (AmBD, Merck UK Ltd, Dorset UK), anti-Mala s 1 mouse monoclonal IgG1 antibody or mouse monoclonal IgG1 antibody isotype control was done by broth microdilution in a 96-well flat-bottomed plate (Fisher Scientific, Leicestershire, UK). The protocol published by the European Committee on Antimicrobial Susceptibility Testing (EUCAST), definite document E.DEF 7.3.1 (Arendrup, 2017) was followed with some modifications. Modifications consisted of using mDixon broth instead of RPMI 1640 and incubation times appropriate for *M. sympodialis* (3 days). Yeasts were grown on agar and in broth as described above. Inoculum was prepared to obtain an absorbance between 0.5-0.1 at 600 nm. Working inocula were obtained by diluting the inoculum 1:10 in mDixon broth and 100 μl of inoculum was added to all wells except for negative controls. Dilutions of each treatment (AmBD, voriconazole, anti-Mala s 1, and IgG1 control) in mDixon broth was added to corresponding wells to give a final volume of 200 μl and concentration of 0.03 to 16 μg/ml. Samples were assayed in triplicate, with a positive control (no drug) and a negative control (medium only) added. Growth was monitored every 6 h over 36 h by measuring optical densities at 530 nm using a microplate reader (VersaMax™ with Softmax Pro 7 software. Molecular Devices LLC, UK) incubating at 37 °C. MICs were also determined by optical density measurements using the same microplate reader at 36 and 96 h of incubation and defined as inhibition of 50% growth for voriconazole or anti-Mala s 1 antibody and 100% growth inhibition for AmBD. These experiments were performed on three biological replicates.

### Skin samples and culture conditions

Human skin tissue without adipose layer from abdominal or breast surgeries was purchased from Tissue solutions^®^ (Glasgow, UK). Skin explants from six different donors, obtained under surgical aseptic conditions, were used to set up a skin model as previously described (Corzo-Leon *et al*, 2021). In brief, the 1 cm^2^ explants were inoculated by applying 1×10^6^ yeast cells in 10 μl directly onto the epidermis. Skin samples with medium only were included as controls in all experiments. Skin samples were recovered after 6 days (1 sample), 7 days (3 samples), 8 days (1 sample), and 14 days (1 sample). Tissue samples for histology were placed into moulds, embedded in OCT Embedding Matrix (Cellpath Ltd, Newtown, UK) and flash-frozen with dry ice and isopentane. These were stored at −70 °C until use. Tissue samples for RNA-protein extraction were cut into small pieces (< 5 mm) and placed in a microcentrifuge tube containing RNAlater^®^ solution (Merck Ltd, Dorset, UK) and stored at −70°C until further use.

### Immunofluorescent staining of skin tissue samples

For histological examination of human explant skin samples (see above), 6 μm skin tissue sections, cut from frozen OCT blocks using a cryostat, were placed on poly-L-lysine coated slides and fixed with 4% methanol-free paraformaldehyde in PBS for 20 min before further staining. After fixation, skin sections slides were washed in ice cold PBS in a Coplin jar. Samples were then permeabilised using 0.2% Triton X-100 for 10 min and washed three times with PBS at room temperature. Slides were incubated in blocking buffer (1% BSA, 0.1% glycine, 0.1% Tween 20 in PBS) for 1 h at room temperature.

Primary antibodies were prepared as described for gold immunolabelling in 1% BSA in PBST (0.1% Tween 20 in PBS). Slides containing 4 sections per slide and 8 sections per sample were incubated with primary antibodies at 4°C overnight and then washed three times with PBS before incubating them with goat anti-mouse IgG (H&L) secondary antibody (Alexa Fluor^®^ 488 in 1% BSA PBST, 1:300; Abcam #ab150113) for 1 h at room temperature in the dark. Samples were then washed three times in PBS. Slides were incubated in 20 μg/ml calcofluor white (Sigma, UK) and 1 μg/ml propidium iodide (HPLC grade; Sigma, UK) for 30 min in a Coplin jar and washed three times with PBS. Slides were left in the dark to air dry. Mounting medium (Vectashield^®^ without DAPI, Vector laboratories Ltd, UK) was added, then a coverslip was added and sealed. Slides were imaged using a Spinning Disk confocal microscope (Zeiss, UK).

### RNA and protein extraction

Skin samples in RNAlater^®^ solution were recovered from storage (see above). RNAlater^®^ solution was discarded and the tissue washed with sterile water. RNA and protein extractions were performed from the same samples in a single two-day process based on previous publications (Chomczynski & Sacchi, 1987, 2006; Berglund *et al*, 2007, Corzo-Leon *et al*, 2019). For *M. sympodialis* yeasts, mDixon broth cultures were harvested by centrifugation, the pellet recovered and resuspended in Trizol for RNA and protein extraction, similar to skin samples. Protein pellets were stored at −20°C in 100 μl re-solubilisation buffer (1% w/v DTT, 2 M thiourea, 7 M urea, 4% w/v CHAPS detergent, 2% v/v carrier ampholytes, 10 mM Pefabloc^®^ SC serine proteinase inhibitor (Thermo-Fisher Scientific, UK)), with protein concentration determined by Coomassie G-250 Bradford protein assay kit following manufacturer’s instructions (Sigma, UK).

### Western Immunoblotting and proteomics

#### SDS-PAGE gel electrophoresis

After protein quantification, 10 μg protein was used for gel electrophoresis and Western Blot. Non-reducing denaturing conditions were used for SDS-PAGE gel electrophoresis. NuPAGE LDS Sample Buffer 4X (Thermo-Fisher Scientific, UK) was added at a 1:3 ratio to protein sample up to a final volume of 10 μl and incubated at 70°C for 10 min. Samples were loaded onto a NuPage 4-12% Bis-Tris gel along with a combination of SeeBlue:Magic Marker (Thermo-Fisher Scientific, UK) 4:1 ratio (final volume 5 μl) and the combination loaded into the first gel lane. The gel was covered in 1X running buffer (Novex NuPAGE MES SDS buffer, Thermo-Fisher Scientific, UK) within an electrophoresis chamber, 500 μl of antioxidant (Invitrogen NuPAGE™ Antioxidant; Thermo-Fisher Scientific, UK) was added to the running buffer covering the surface of the gel. Electrophoresis was carried out for 40-55 min at 150 V.

#### Protein membrane transfer

Proteins were transferred to a PVDF membrane (Thermo-Fisher Scientific, UK) using ice cold 1X transfer buffer (NuPAGE transfer buffer, 10% methanol in deionised water). The PVDF membrane was activated prior to transfer in 100% methanol for 1-2 min, distilled water and transfer buffer. Transfer was carried out at 25 V for 2 h 15 min. The chamber was kept on ice during the transfer process.

#### Western blots

PVDF membrane were retrieved after transfer and washed once with distilled water. The membrane was blocked at room temperature for 1 h in blocking buffer (5% skim milk in 0.1% PBS-Tween 20). The membrane was incubated overnight with the primary antibody (1:1000 in 5% BSA in 0.1% PBS-Tween 20) at 4°C in 50 ml plastic tubes on a tube roller. The membrane was washed five times for 5 min with 0.1% PBS-Tween 20 before being incubated for one h at room temperature with goat antimouse IgG H&L secondary antibody (1:2000 in 5% skim milk in 0.1% PBS-Tween 20). Finally, membranes were washed five times for 5 min with 0.1% PBS-Tween 20 before being treated with SuperSignal^TM^ West Femto (Thermo-Fisher Scientific, UK). The chemiluminescent signal was captured with the Peqlab camera (VWR, UK) and analysed using Fusion molecular imaging software v 15.5 (Fisher Scientific, UK).

#### Proteomics and bioinformatics analysis

Western blots (as described above) of *M. sympodialis* proteins and proteins from human skin were performed to determine cross-reactivity of the Mala s 1 antibody. The gel was stained with Coomassie Brilliant Blue (CBB) G250 (Thermo-Fisher Scientific, UK) and bands cut out for automated in-gel digestion and identification by Liquid Chromatography with tandem mass spectrometry (LC-MS/MS) using a Q Exactive Plus (Thermo Fisher Scientific, UK) at the University of Aberdeen proteomics facility.

Gel band proteins identified by mass-spectrometry were analysed by applying several filters. The first filter applied selected proteins by the expected protein size, based on the predicted kDa of the band detected by the anti-Mala s 1 antibody on the western blot. Then, only proteins with two or more identified peptides and two or more peptide-spectrum matches (PSM) were selected. Finally, proteins were selected by cellular localisation using the Uniprot database (https://www.uniprot.org/).

#### DNA extraction, ITS PCR and pGEM-T vector cloning

Genomic DNA was extracted (Hoffman & Winston, 1987) from control skin explants (Corzo-Leon *et al*, 2020) or *M. sympodialis*-inoculated skin explants stored in OCT Embedding Matrix. As a positive control, DNA was extracted from *Candida albicans* (ATCC 90028) pure culture grown in YPD broth (1% w/v yeast extract, 2% w/v peptone, 2% w/v glucose). Distilled water was included as a negative control.

To identify any fungi in the skin samples, the ITS1 primer pair (ITS1F (5’CTTGGTCATTTAGAGGAAGTAA 3’), ITS1R (5’ GCTGCGTTCTTCATCGATGC 3’ (Irinyi *et al*, 2015b, 2015a)) were used. The ITS1 region was amplified using 2X KAPA HiFi Hotstart ReadyMix following manufacturer’s recommendations (Roche, UK), with 0.4 μM primers, and 50 ng template DNA. PCR settings were: one cycle of initial denaturation at 95 °C for 3 min, 30 cycles of denaturation at 98 °C for 20 s, annealing at 52 °C for 15 s, extension at 72 °C for 30 s and one cycle of final extension 72 °C for 1 min. PCR products were analysed on a 1% agarose TAE gel.

PCR products were purified with PEG/NaCl purification buffer for 60 min at room temperature followed by centrifugation for 1 h at 2,200 x g at 4 °C. The supernatant was removed and DNA precipitated in 70% ethanol, and the DNA pellets were recovered with centrifugation at 2,200 x g for 10 min at 4 °C and resuspended in sterile water. A-tailing of purified PCR products was performed using GoTaq polymerase (Promega, UK) according to the manufacturer’s instructions. The A-tailed PCR products were ligated into Promega pGEM-T vector following the manufacturer’s instructions. Plasmid extraction was carried out with Qiagen QIApep^®^ spin miniprep kit following manufacturer’s instructions (QIAGEN Ltd, UK). Inserts were confirmed by EcoRI restriction digestion and gel electrophoresis. The recovered plasmids sequenced by DNA Sequencing & Services (MRC I PPU, School of Life Sciences, University of Dundee, Scotland). Sequences were analysed using SeqMan Pro (DNASTAR, Madison, WI US), and species identified by BLAST (https://blast.ncbi.nlm.nih.gov/).

#### In silico *sequence analysis of Mala s 1*

The Mala s 1 protein sequence (ID M5E589) was downloaded from Uniprot (www.uniprot.org/). Presence of a KELCH domain in the Mala s 1 protein sequence was investigated using a previously published consensus KELCH domain sequence (Adams *et al*, 2000; Prag & Adams, 2003). The potential epitope bound by the anti-Mala s 1 antibody was manually identified in the Mala s 1 protein sequence based on previous publications (Schmidt *et al*, 1997; Gioti *et al*, 2013).

## Results

### *Mala s 1 is present in the cell wall of* M. sympodialis

*M. sympodialis* yeast cells were processed by HPF and freeze substitution to localise Mala s 1 by TEM and immunohistochemistry. Gold immunolabelling of yeast cells incubated with the anti-Mala s 1 antibody showed that Mala s 1 was found mainly in the cell wall and in the budding area (Figure 1). Gold particles bound to the Mala s 1 allergen were grouped in pairs, triplets and quadruplets and binding was not specific to a particular region of the cell wall. Some gold labelling was also detected inside the yeast cells but mainly close to the cell wall rather than in the centre of the cell. No gold particles were observed either intracellularly or bound to the cell walls when yeast cells were incubated with the gold conjugated secondary antibody only or the control IgG1 antibody (Supplementary Figure 1).

**Figure 1.**
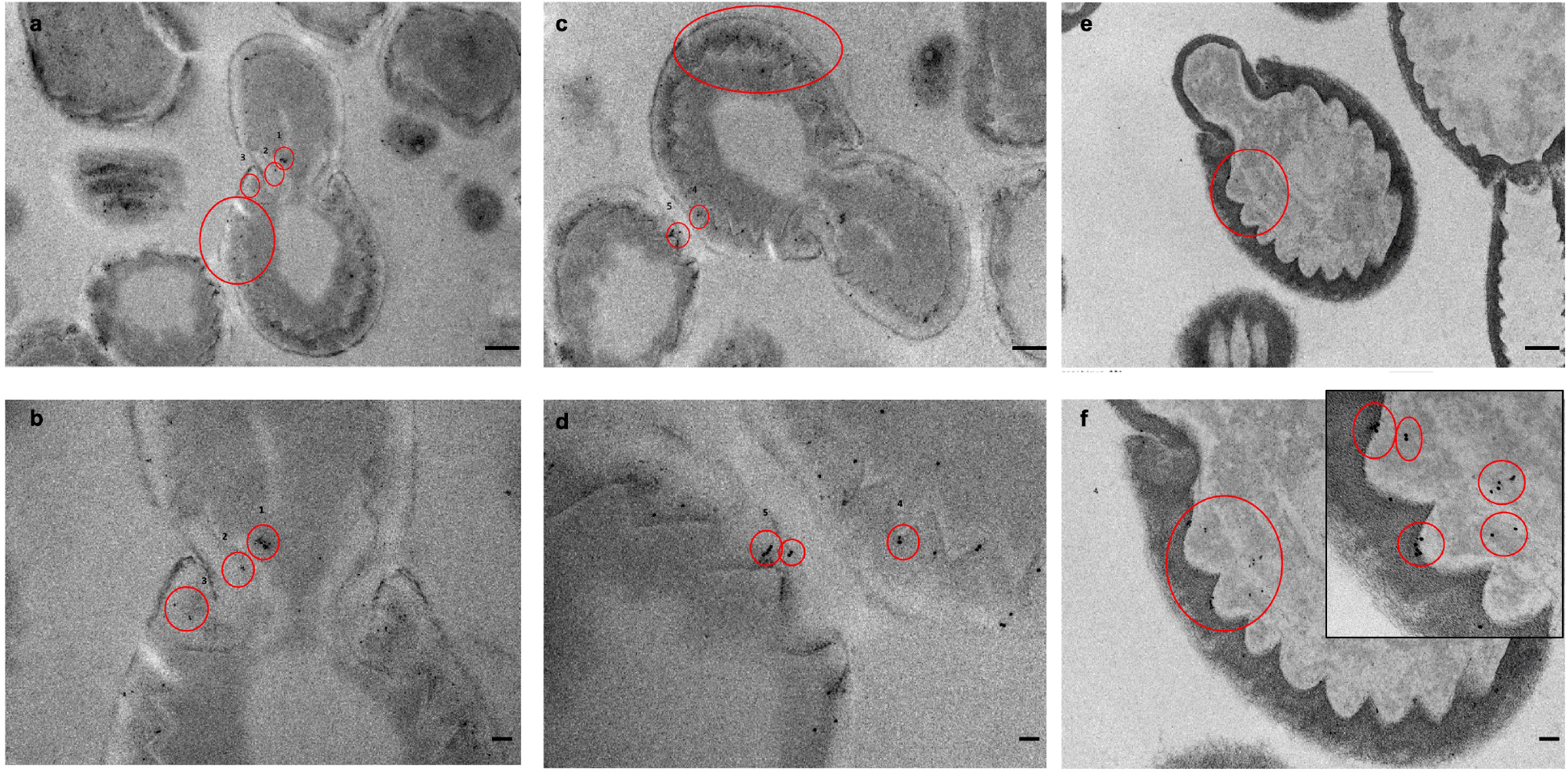
Localisation of Mala s 1 on *M. sympodialis* yeast cells. TEM images of *M. sympodialis* yeast cells cultured on mDixon agar for 4-5 days stained by gold immunolabelling with the anti-Mala s 1 antibody and a secondary gold conjugated antibody. Red circles indicate colloidal gold particles often grouped in pairs, triplets and quadruplets. (a,b) The same yeast cell at different magnification showing Mala s 1 at the budding site. (c,d) Two yeast cells at different magnification showing Mala s 1 in the cell wall. (e,f) A yeast cell showing Mala s 1 close to the cell wall at three different magnifications. Scale bars in images: (a, c, e) = 500 nm, (b, d, f) = 100 nm, and insert in (f) = 50 nm. Images are representative of duplicate experiments.

### Mala s 1 is not an antifungal target

We then assessed whether co-culture of *M. sympodialis* cells with the anti-Mala s 1 antibody could interfere with yeast growth using a modified EUCAST broth microdilution antifungal susceptibility assay (Arendrup *et al*, 2017). Amphotericin B deoxycholate (AmBD) and voriconazole were used as positive antifungal controls and a non-specific IgG1 antibody used as a negative control, with concentrations ranging from 0.03 to 16 μg/ml. During the first 36 h of incubation, anti-Mala s 1 antibody-treated yeasts grew slower than untreated yeasts (Figure 2a), however this same pattern was observed for yeasts treated with the IgG1 control, as well as voriconazole and AmBD (Figure 2b–2d). After 36 h, all concentrations of voriconazole and AmBD inhibited growth by at least 50% (Fig 2e). After 96 h, only voriconazole had more than 80% inhibitory activity at all concentrations tested (Figure 2f). After 96 h, samples were cultured on mDixon agar without drug to identify the minimal effective, or minimum fungicidal concentration (See Suppl Fig. 2). Only voriconazole was fungicidal for *M. sympodialis* at all concentrations tested (from 0.01 μg/ml up to 8 μg/ml) (Supplementary Figure 2).

**Figure 2.**
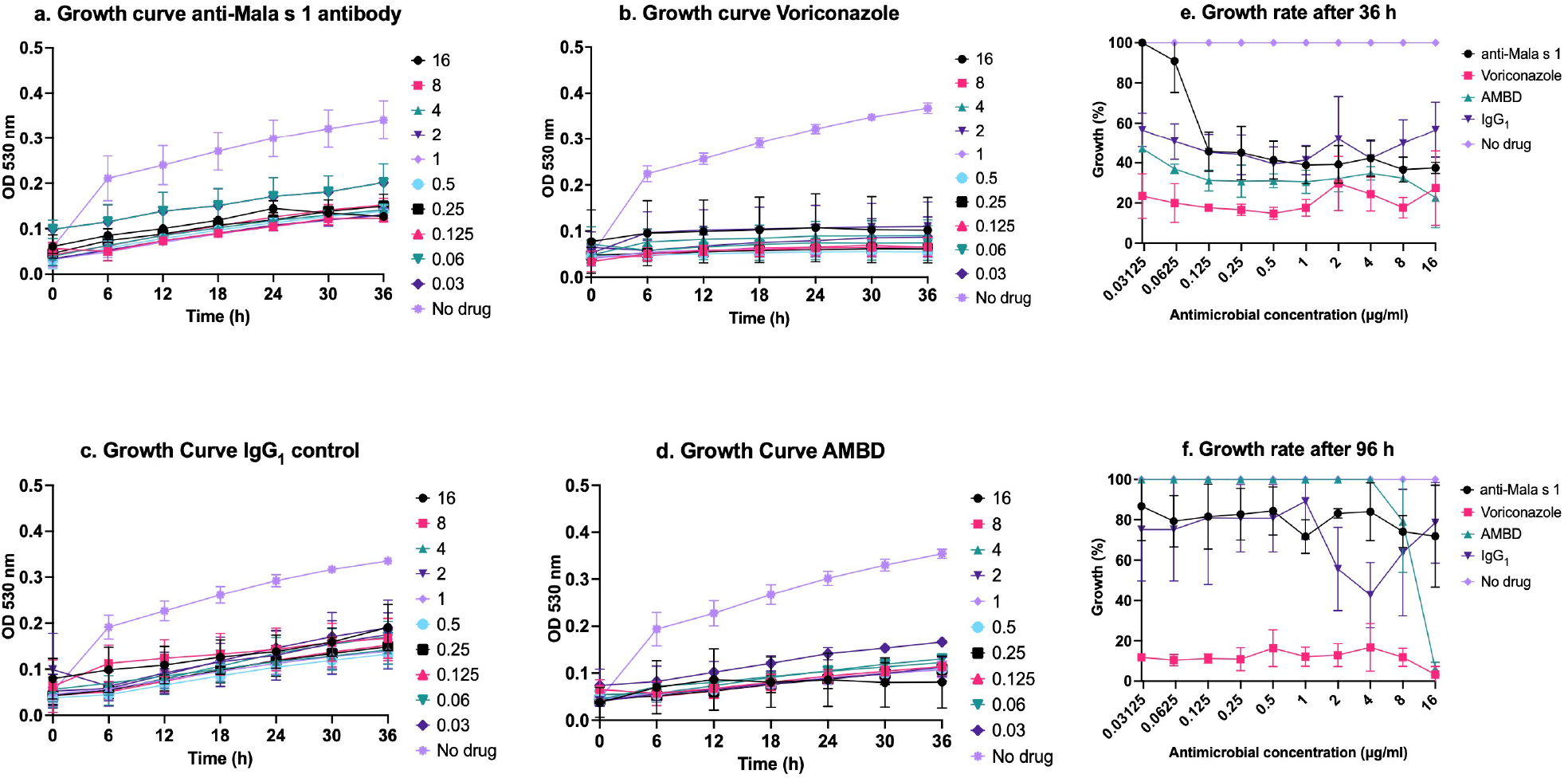
Antifungal activity of the anti-Mala s 1 antibody, AmBD or voriconazole against *M. sympodialis* yeasts. (a-d) 36 h growth curves presenting average ODs measured every 6 h at 530 nm. *M. sympodialis* was co-cultured with (a) anti-Mala s 1 antibody, (b) voriconazole, (c) IgG control, and (d) AmBD. Growth curves represent the mean of three biological replicates, with each colour representing a different drug concentration, ranging from 16 to 0.03 μg/ml or untreated yeasts. Growth was plotted after 36 h (e) and 96 h (f) as percentage of growth when compared to no drug control. Plotted colour lines correspond to the mean of three biological replicates, error bars correspond to standard deviation and different colours correspond to different drugs or no drug control.

### Anti-Mala s 1 antibody cross-reacts with proteins in human non-inoculated skin

Next, we investigated if the anti-Malas s 1 antibody recognised human skin proteins present in skin explants as indirect evidence of potential cross-reactivity between Mala s 1 and human skin proteins. Non-inoculated tissue from six different skin donors was analysed by immunofluorescence microscopy using the anti-Mala s 1 antibody. For all donors, samples stained with the anti-Mala s 1 antibody showed detectable fluorescent signal when compared to controls (only secondary antibody or IgG1 isotype control) (Figure 3). The anti-Mala s 1 antibody stained the epidermis (as shown by colocalisation with propidium iodide-stained epidermal cells) with stronger signals in the basal layers, intercellular junctions, as well as keratinocyte cytoplasm and nucleus (Supplementary Figure 3). There was some variation in staining between different donors (Supplementary Figure 3). Calcofluor white staining was used to indicate the presence of fungal structures on the skin. No chitin was detected, indicating that no fungi were present on the skin explants (Figure 3).

**Figure 3.**
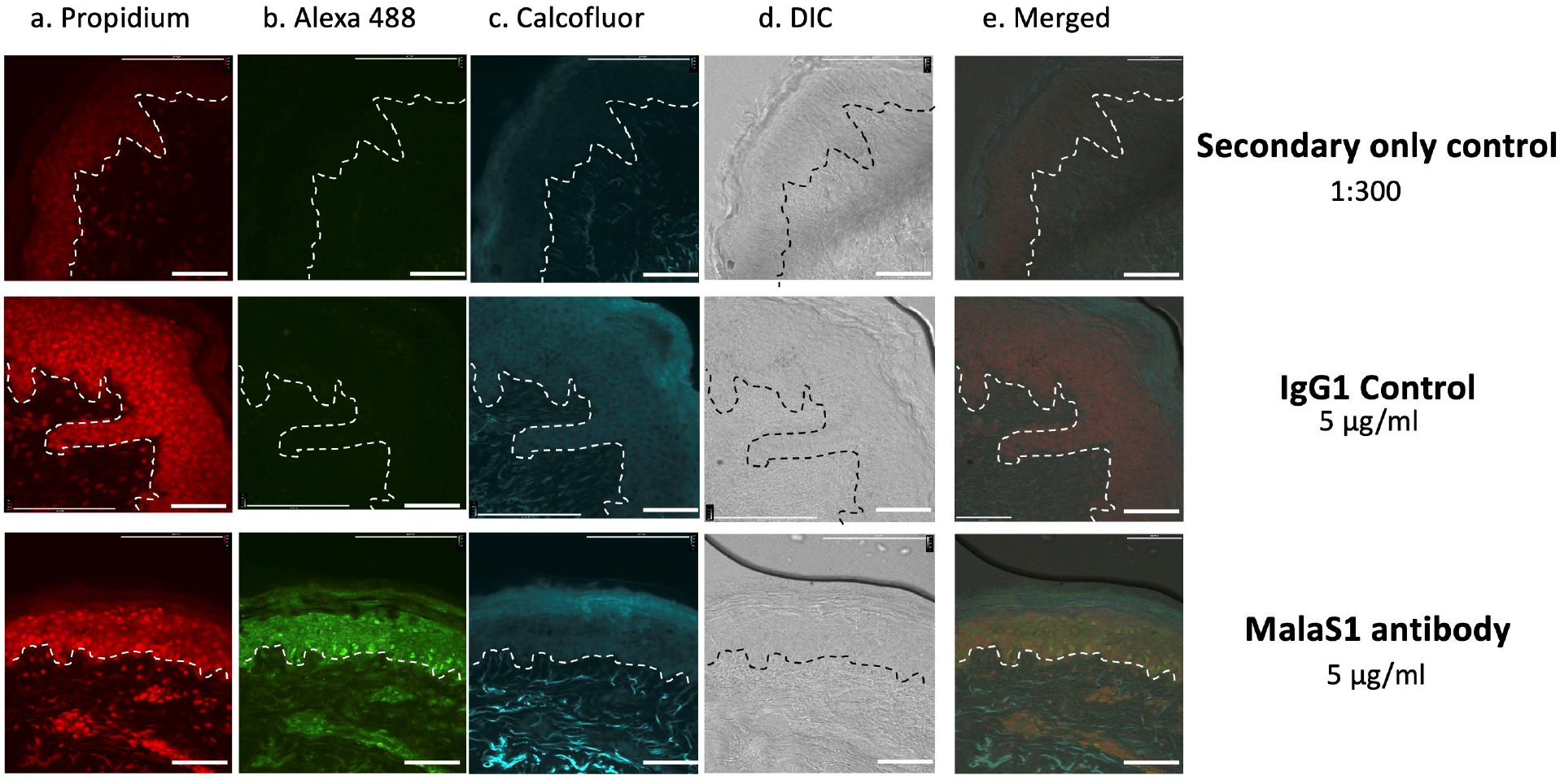
Anti-Mala s 1 antibody reacts with human proteins in non-inoculated skin. Sections were imaged with (a) Rh-TRITC channel (543 nm/569 nm, propidium iodide staining), (b) FITC channel (495 nm/519 nm, stained with secondary antibody labelled with Alexa 488), (c) DAPI channel (358 nm/461 nm, calcofluor staining), (d) images captured with differential interference contrast (DIC), (e) merge of all channels. 6 μm thin tissue sections were cut from frozen OCT skin sample blocks. Samples were stained with the Mala s 1 antibody and compared to controls using only the secondary antibody labelled with Alexa 488, and a mouse IgG1 control. Images are representative of skin samples from six different donors. White line separates epidermis (top) from dermis (bottom). Scale bar is 100 μm

These non-inoculated skin samples were checked for the presence of *Malassezia* DNA to rule out prior colonisation of the skin samples by *Malassezia* species or contamination of the skin during experiments. The ITS1 region was amplified and sequenced from DNA extracted from the six noninoculated skin samples and compared with control skin samples previously inoculated with *M. sympodialis* and *C. albicans*. Four out of the six non-inoculated skin samples amplified ITS1 PCR products, but no *M. sympodialis* sequences were identified in any of the six non-inoculated samples (Supplementary Figure 4). One skin sample (4-17) contained a sequence of *Malassezia globosa* (Supplementary Figure 4). Sequence analysis identified *C. albicans* and *M. sympodialis* from inoculated skin samples, as expected.

Proteins were then extracted from non-inoculated human skin from three donors to detect potential anti-Mala s 1 antibody targets in skin. Proteins were also extracted from *M. sympodialis* yeast cells grown on mDixon agar and in mDixon broth as Mala s 1 protein controls. Expression of the Mala s 1 protein was evaluated by immunoblotting. The anti-Mala s 1 antibody correctly identified a 37 kDa protein corresponding to the size of the Mala s 1 protein in *M. sympodialis* yeast cells grown on mDixon agar (Figure 4a) and in mDixon broth (data not shown). The anti-Mala s 1 antibody also identified a protein ≥200 kDa in protein extracts from non-inoculated skin samples (Figure 4b). These gel bands were recovered and processed by automated in-gel trypsin digestion and the peptides separated and identified by LC-MS/MS. To identify the potential protein(s) recognised by the Mala s 1 antibody, proteins identified by proteomics (n=180) were filtered, initially by selecting proteins of expected size (≥ 200 kDa= 17 proteins found), then proteins where the number of peptides were ≥2 and peptide spectrum matches (PSM) was ≥2 (n=12). Next, proteins were filtered according to their cellular localisation and protein structure (n=6). MalaEx can also contain Mala s 1 and can be taken up by human keratinocytes and monocytes where MalaEx are mainly found in close proximity to the nuclei (Johansson *et al*, 2018). Therefore, proteins localising to the cell nucleus were also included. Considering the antibody binding pattern observed in skin samples (Figure 3), an extra filter was applied for proteins expressed in all epidermal layers and in nuclei, cytoskeleton, and cytoplasm. After applying all filters, two proteins remained as potential targets: epiplakin and desmoplakin (Figure 4c).

**Figure 4.**
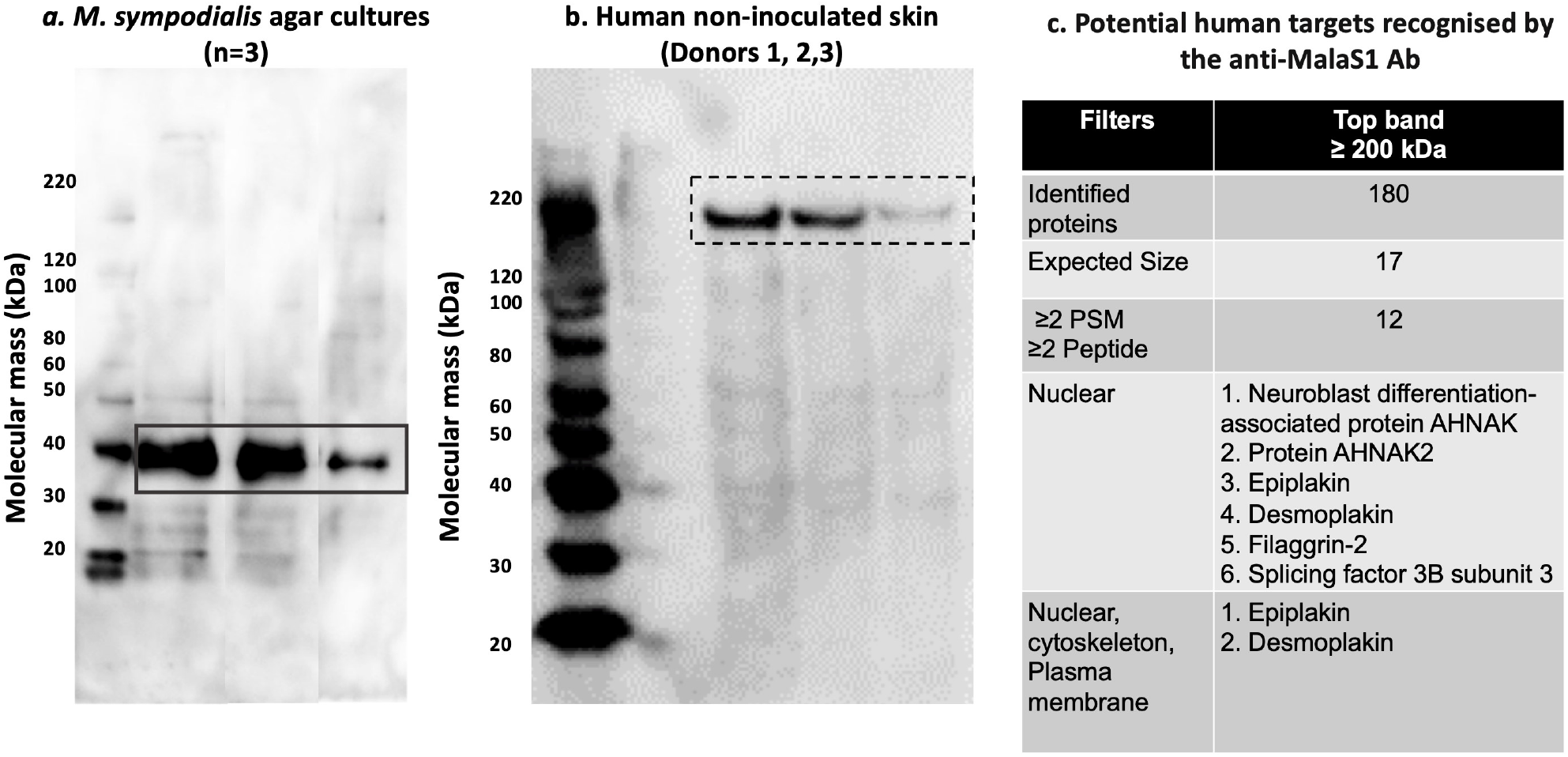
Mala s 1 expression in *M. sympodialis* yeast cells and in human skin. Image (a) Protein detected by the anti-Mala s 1 antibody in three different *M. sympodialis* cultures on mDixon agar for 4-5 days and (b) in human non-inoculated skin samples from three different donors). Immunoblotting was carried out under denaturing, non-reducing conditions using the anti-Mala s 1 primary antibody. In (a) black continuous-line squares surround the Mala s 1 expected size (37 kDa) and in part (b) black dotted-line squares surround high molecular protein bands recognised by the Mala s 1 antibody. (c) shows potential human proteins recognised by the anti-Mala s 1 antibody after protein sequencing and filtering.

### Mala s 1 is a potential KELCH protein

Mala s 1 is a β-propeller protein (Vilhelmsson *et al*, 2007b) containing 4-8 symmetrical beta sheets. We show here that the anti-Mala s 1 antibody-stained human skin sections and previous data indicated that MalaEx vesicles were taken up intracellularly in keratinocytes and monocytes (Johansson *et al*, 2018). This led us to test the hypothesis that Mala s 1 might be a β-propeller protein with a KELCH motif and hence act as an allergen due to molecular mimicry with human skin proteins. The 350 aa Mala s 1 protein sequence (Uniport ID M5E589) was analysed manually for KELCH motifs using the previously published motif consensus (Prag & Adams, 2003; Adams *et al*, 2000). A 64 aa sequence was identified as having the consensus for a KELCH motif (Figure 5). The anti-Mala s 1 antibody is known to recognise the peptide epitope (SFNFADQSS) (Schmidt *et al*, 1997) which lies out with the predicted KELCH motif of Mala s 1, but no sequences resembling the antibody target peptide were detected in the two potential human target proteins, epiplakin and desmoplakin. The antibody target epitope sequence was only found in the Mala s 1 protein sequence (Figure 5).

**Figure 5.**
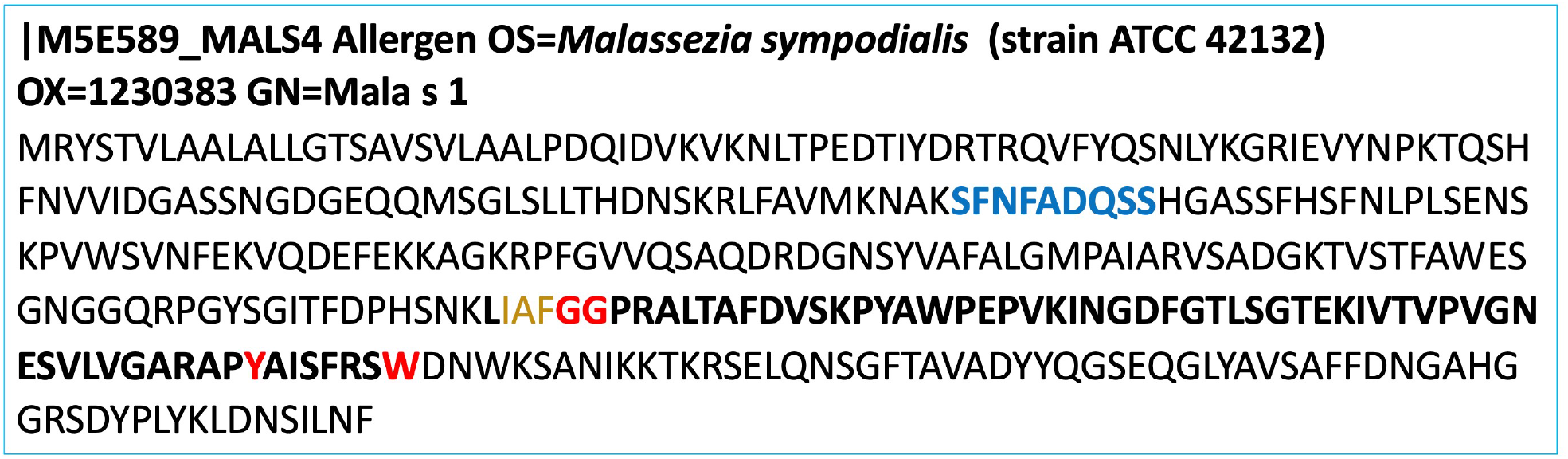
Mala s 1 is a potential KELCH protein. Mala s 1 sequences obtained from Uniprot (M5E589). *In silico* analysis of Mala s 1 identified a KELCH motif (highlighted in bold). Mala s 1 has a β-propeller structure (Vilhelmsson *et al*, 2007b), and three hydrophobic amino acids (brown) are found before a double GG (red font), and six aa residues are located between Y (red font) and W (red font). The blue font indicates the peptide sequence reported as the epitope of anti-Mala s 1 antibody (Schmidt *et al*, 1997).

## Discussion

Immunostaining with TEM confirmed that Mala s 1 is localised in the *M. sympodialis* cell wall. Binding of the anti-Mala s 1 specific antibody to its cell wall target did not inhibit growth of the fungus suggesting that Mala s 1 is not an antifungal therapeutic target. The anti-Mala s 1 antibody recognised and bound to unknown targets in full thickness human skin explants. This cross-reactivity could be due to anti-Mala s 1 antibody recognising human proteins with similar tertiary structures or epitopes to Mala s 1 despite Mala s 1 not having a clear human homologue. Proteomic analysis of anti-Mala s 1 antibody-reactive proteins isolated from western blotting revealed a number of candidate human targets, but none contained the specific peptide epitope recognised by the antibody on Mala s 1. Bioinformatics analysis predicted that Mala s 1 contains a KELCH domain, a common motif for protein-protein interactions. This poses the question whether the Mala s 1 KELCH domain is responsible for cross-reactivity between *anti-Malassezia* antibodies and human skin proteins.

Taking advantage of an anti-Mala s 1 specific mouse monoclonal antibody (Zagari *et al*, 1994, Schmidt *et al*, 1997), we demonstrated by TEM immuno-gold labelling that Mala s 1 is mainly localised to the cell wall of *M. sympodialis* yeast cells, in agreement with Zargari *et al*, (1997). The immuno-gold labelling was found in a disperse pattern around the cell periphery of the mother cell wall, but also in the budding area and inside yeast cells (Figure 1). As immuno-gold labelled Mala s 1 particles appeared to be enriched at the plasma membrane/cell wall interface we cannot rule out that we are also visualizing the MalaEx vesicles (Gehrmann *et al*, 2011), which are enriched in Mala s 1 (Johansson *et al*, 2018).

Although the anti-Mala s 1 antibody recognised and bound the *M. sympodialis* yeast cell wall, drug susceptibility testing did not show an antifungal effect against *M. sympodialis* (Figure 2). This suggested that Mala s 1 is not essential for replication nor as a cell wall component; hence, Mala s 1 is not a potential target for antifungal therapy. AmBD was also ineffective in the drug susceptibility testing performed in this study, whereas voriconazole was fungicidal at low drug concentrations. Previous studies have documented higher susceptibility of *Malassezia* species to triazoles, specifically triazoles with anti-mould activity (Rojas *et al*, 2014; Miranda *et al*, 2007; Carrillo-Muñoz *et al*, 2013; Leong *et al*, 2017). Contrary to this study, AmBD has been reported to achieve 100% growth inhibition at high MICs (ranging from 0.5 to 4 μg/ml) for cutaneous clinical isolates (Rojas *et al*, 2017, 2014; Leong *et al*, 2017) and MICs ≥ 8 μg/ml to AmBD were documented for blood-recovered *Malassezia furfur* (Iatta *et al*, 2014). These differences could be explained by the type of medium used for antifungal susceptibility testing, mDixon vs supplemented RMPI 1640 or the fungal isolates used in this study. In the future the anti-Mala s 1 antibody used in combination with antifungal drugs could be tested for synergy of the antibody with other antifungal therapies. Potential beneficial immunomodulatory effects and effector functions of the anti-Mala s 1 antibody could also be tested in assays with immune cells or in other *ex vivo* human-pathogen interaction models.

Since we found that the anti-Mala s 1 antibody did recognise human proteins present in noninoculated human skin explants, we used a combination of bioinformatics and proteomics to investigate potential human skin orthologous proteins that may contribute to the host’s inflammatory responses to this allergen. Although several candidate proteins were identified, such as epiplakin and desmoplakin (Figure 4), we cannot definitively pinpoint that the anti-Mala s 1 antibody cross reacts with a single human skin protein. Bioinformatics analysis revealed for the first time that Mala s 1 allergen is a potential KELCH protein. Proteins with KELCH motifs have wide-ranging functions including cellular communication, with examples that contribute to microbial pathogenicity and skin function relevant to this study. Two important examples of these KELCH proteins are the human intracellular NS1-binding protein (NS1-BP) targeted by NS1 protein of Influenza A virus during infection (Wolff *et al*, 1998) and the KELCH K13-propeller protein in *Plasmodium falciparum* where SNPs are associated with artemisinin resistance, which limits the eradication of malaria in areas where these mutations are highly prevalent, for example Southeast Asia and China (Ménard *et al*, 2016). Another example of a KELCH protein complex that functions specifically in the skin is the nuclear factor erythroid-2-related factor 2/KELCH-like ECH-associated protein 1 (Nfr2/KEAP1) complex (Helou *et al*, 2019). In this complex, Nfr2 initiates the transcription of antioxidant enzymes, limits both the levels of reactive oxygen species (ROS) and inflammatory responses, such as inflammasomes, while KEAP1 supresses these functions (Helou *et al*, 2019). Dysregulation of the Nfr2/KEAP1 system has been associated with skin damage and psoriatic plaque formation in a psoriasis-mouse model (Ogawa *et al*, 2020). Considering the multiple functions of KELCH proteins in the skin and their roles in the pathogenesis of human infections, the study of Mala s 1 allergen and determining its role during interaction with skin is highly relevant.

In the current study, two potential human target proteins that cross-react with Mala s 1 antibody were identified: epiplakin and desmoplakin (Figure 4). The distribution of these two proteins in the epidermis was compared to the observed binding pattern of anti-Mala s 1 antibody to human skin. Desmoplakin (260 kDa) is closer to the size of the immunoblot band than epiplakin (460 kDa).

Desmoplakin has a Src homology 3 (SH3) domain containing 6 β-strands in 2 anti-parallel β-sheets that is key in substrate recognition, membrane localisation and regulation of kinase activity (Kurochkina *et al*, 2013). Desmoplakin contains the 8 highly conserved residues, predictive of a KELCH motif consensus sequence between residues 480 and 540. Four of these residues are localised within the SH3 domain. SH3 domains can interact with KELCH proteins. For example, SH3 domain in Tyrosine-protein kinase Fyn has been described as interacting with KLHL2 (KELCH Like Family Member 2) which has an important role in ubiquitination and actin cytoskeleton formation (Dhanoa *et al*, 2013). Desmoplakin is essential in forming desmosomes, which are cell-cell adhesions necessary for skin integrity (Vasioukhin *et al*, 2001). Recessive mutations in *DSP* (the desmoplakin gene) have been associated with palmoplantar keratoderma, acantholytic epidermolysis bullosa, dermatitis and extensive skin erosion (McAleer *et al*, 2015).

In conclusion, we propose that the IgE binding allergen Mala s 1 of *M. sympodialis* is a potential KELCH protein with a motif that is found in a diverse set of proteins, including over 100 human proteins, which can fold into a β -propeller. We suggest that the similarity of Mala s 1 to human proteins may lead to cross-reactivity and thereby contribute to the skin inflammation amongst AE patients sensitized to *M. sympodialis*.

## Acknowledgments

We thank Giuseppe Ianiri and Joe Heitman for their continuous support and many insightful discussions. Thanks to the Microscopy and Histology Facility at the Institute of Medical Sciences, University of Aberdeen, for sample processing and access to microscopes. Thanks to Dr David Stead and the Aberdeen Proteomics Facility, University of Aberdeen for the proteomics analysis.

## Author Contributions Statement

C. M., A.S., and D.M. conceived the idea and designed the study. D.E.C.L contributed to experimental design and performed the experiments. C.M. and D.M. contributed to project management, and supervision. All authors participated in data interpretation. D.E.C.L. drafted the first version of the manuscript. All authors contributed in writing, critically reviewed and edited the manuscript, and approved the final version.

## Supplementary material

**Supplementary Figure 1. TEM images of *M. sympodialis* yeast cells cultured in mDixon agar for 4-5 days. Images correspond to negative controls for gold immunolabelling**. Yeasts were stained using gold conjugated secondary antibody only (a,b) and IgG1 isotype control (c,d). Scale bars in (a, c)= 500 nm and in (b, d) = 100 nm. Images are representative of 2 experiments.

**Supplementary Figure 2. Minimal effective concentration (MEC) for the anti-Mala s 1 antibody, AmBD, voriconazole and isotype IgG1 control.** Five μl samples were removed from mDixon broth microdilution assays after 96 h of incubation and were cultured on mDixon agar. (a) Each spot represents growth of viable cells after incubation in different drug concentrations ranging from 8 to 0.03 μg/ml in the MIC tests. (b) For the IgG1 control representative values from 1 to 0.06 μg/ml are shown. Experiments were performed in triplicate, one representative presented.

**Supplementary Figure 3. Anti-Mala s 1 antibody reacts with human proteins in non-inoculated skin from 3 donors.** Images of skin samples from three different donors stained with propidium iodide, anti-Mala s 1 antibody and secondary antibody labelled with Alexa 488 and merged channels.

**Supplementary Figure 4. ITS1 PCR amplification products and sequencing results from noninoculated human skin samples.** (a) ITS1 PCR amplification products from DNA extracted from six different skin donors. Lane 1 100 bp ladder, lanes 2-7 are non-inoculated skin samples (labelled month/year of the experiment), lane 8 skin sample inoculated with *M. sympodialis* yeasts, Lane 9 is water and lane 10 are ITS1 amplification products from DNA of *C. albicans* ATCC 90028. (b) Sequencing results of the ITS1 PCR amplification products corresponding to each lane in gel.

