## Supplemental Figures for "*Malassezia sympodialis* Mala s 1 allergen is a potential KELCH protein that cross reacts with human skin"

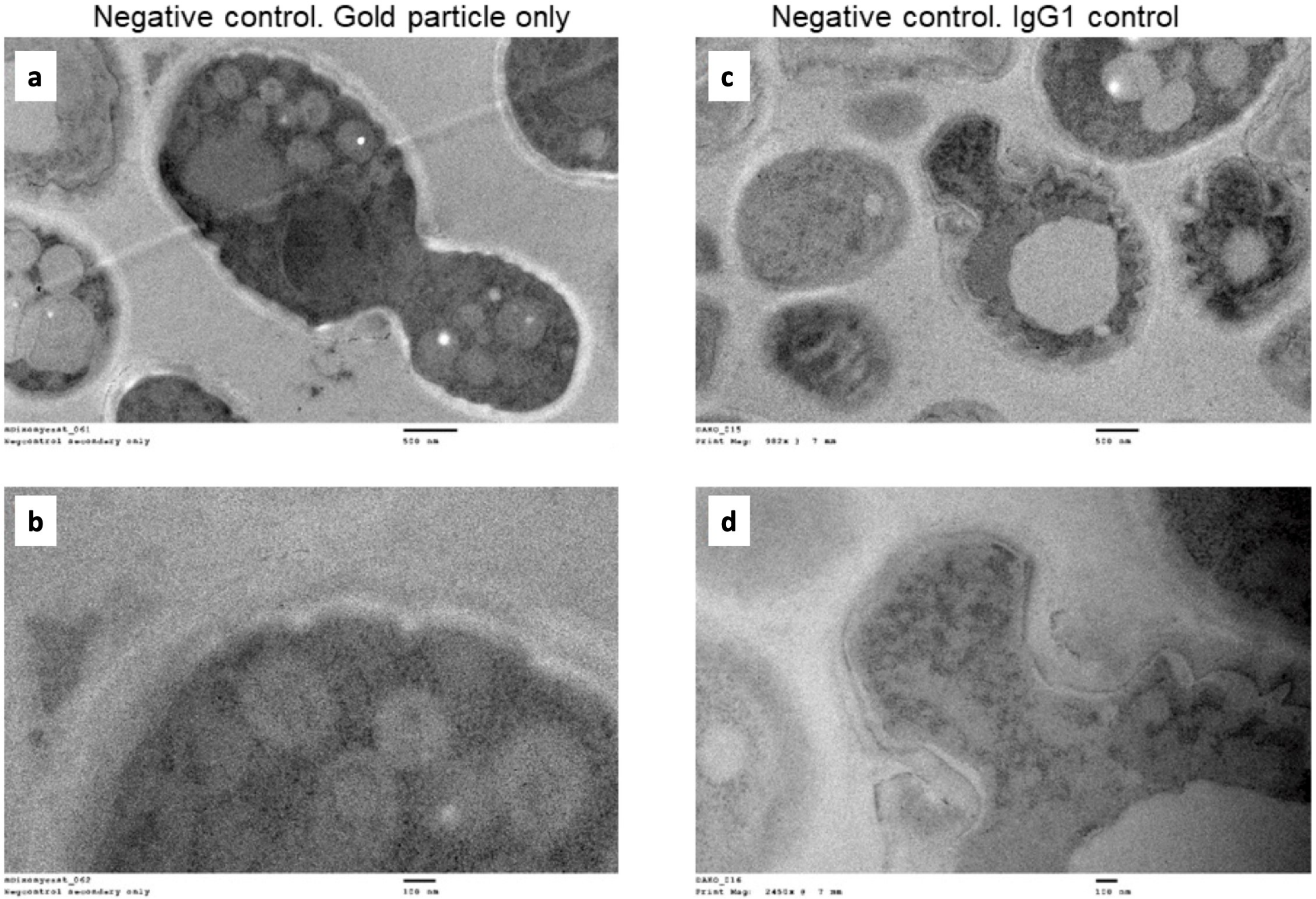
Negative control. Gold particle only
Negative control. IgG1 control
a
b
c
d

**Figure S1**

### Minimal effective concentration after 96 h of incubation

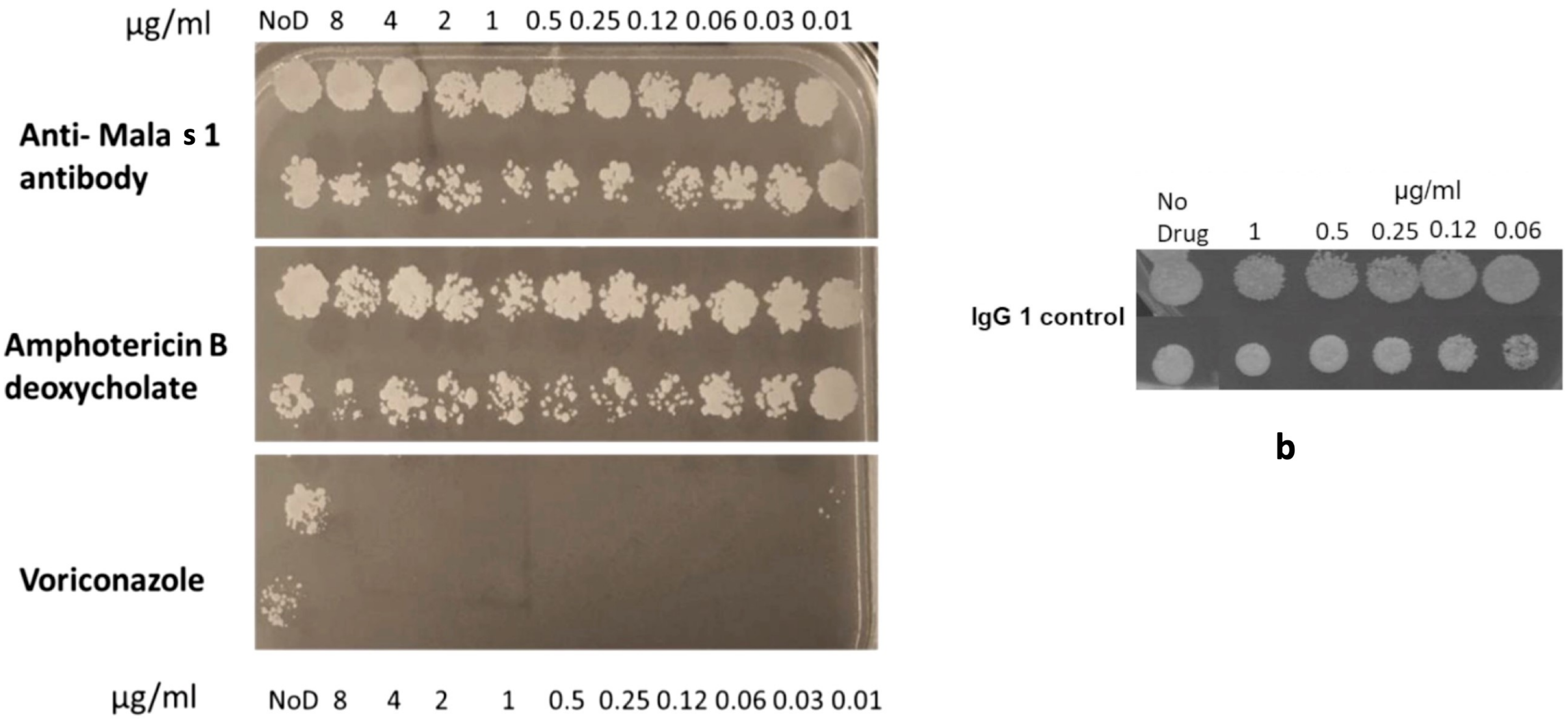


a

b

Figure S2

#### Non-infected human skin (donors 1, 2, 4)

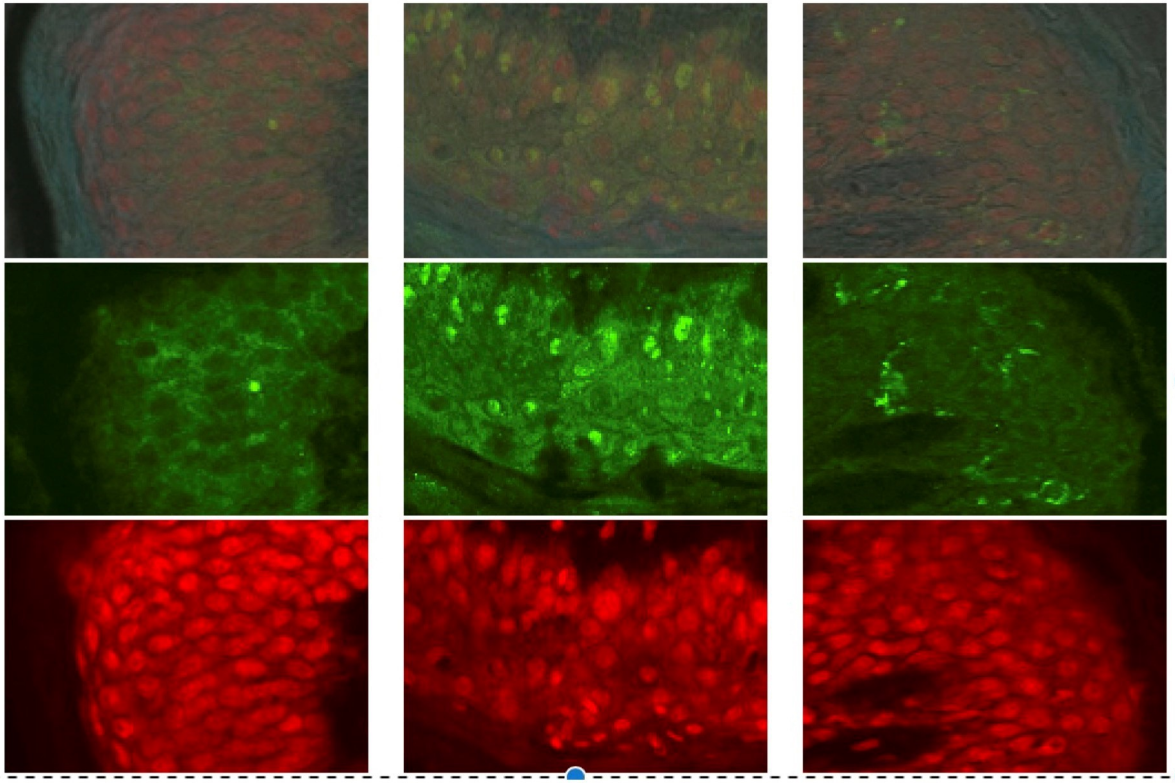


**Figure S3**

**a**

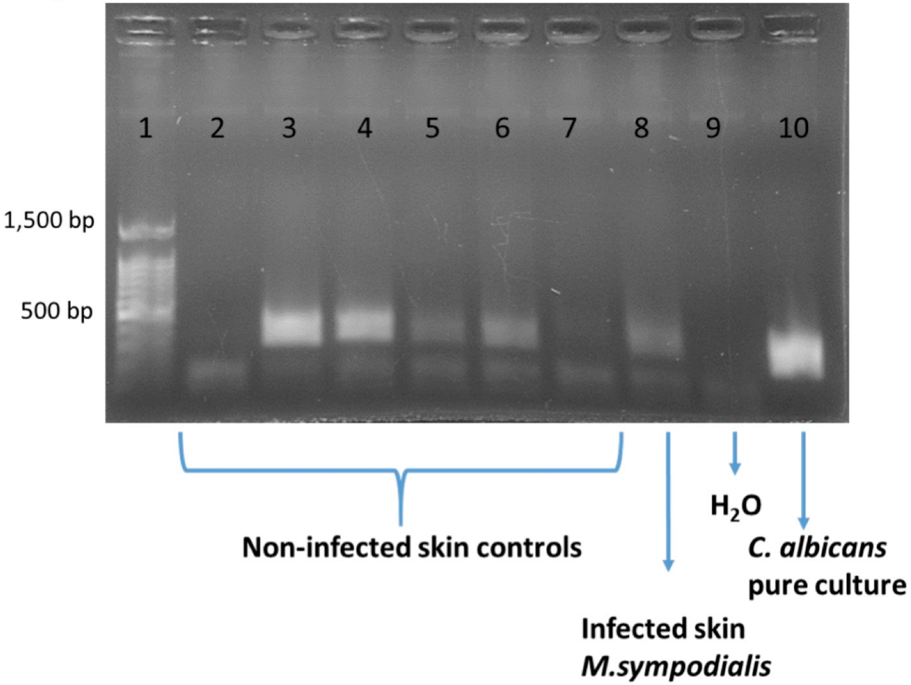


**b**

| Lane | Sample | Amplified | ID in sequencing |
|---|---|---|---|
| 1 | 100 bp ladder | | |
| 2 | Uninfected skin (3-18) | NO | |
| 3 | Uninfected skin (11-17) | YES | 2/5 *Sarocladium kiliense*<br>3/5 Uncultured fungi |
| 4 | Uninfected skin (4-17) | YES | 1/5 *M. globosa*,<br>2/5 *S. schenckii*,<br>2/5 *Sarocladium kiliense* |
| 5 | Uninfected (1-18) | YES | 3/5 *C. albicans*<br>2/5 no inserts |
| 6 | Uninfected skin (6-18) | YES | 2 plasmids with unspecific sequence inserts |
| 7 | Uninfected skin (5-17) | NO | |
| 8 | *M. sympodialis* Infected skin | Yes | 2 plasmids with *M. sympodialis* inserts |
| 9 | Water | No | |
| 10 | *C. albicans* ATCC 90028 | Yes | *C. albicans*, 3/3 plasmids |

**Figure S4**
